## Supplementary data for "Projections from thalamic nucleus reuniens to hippocampal CA1 area participate in context fear extinction by affecting extinction-induced molecular remodeling of excitatory synapses"

[#] currently Electron Microscopy Platform & Bio Imaging Core at VIB-KU Leuven Center for Brain and Disease Research, and Department of Neurosciences at KU Leuven, Campus Gasthuisberg, building O&N5, Herestraat 49 box 602, 3000 Leuven, Belgium.

***Corresponding author:** Kasia Radwanska, Ph.D., Laboratory of Molecular Basis of Behavior, the Nencki Institute of Experimental Biology of Polish Academy of Sciences, 3 Pasteur St., Warsaw 02-093, Poland;; tel: +48 501 736 942.

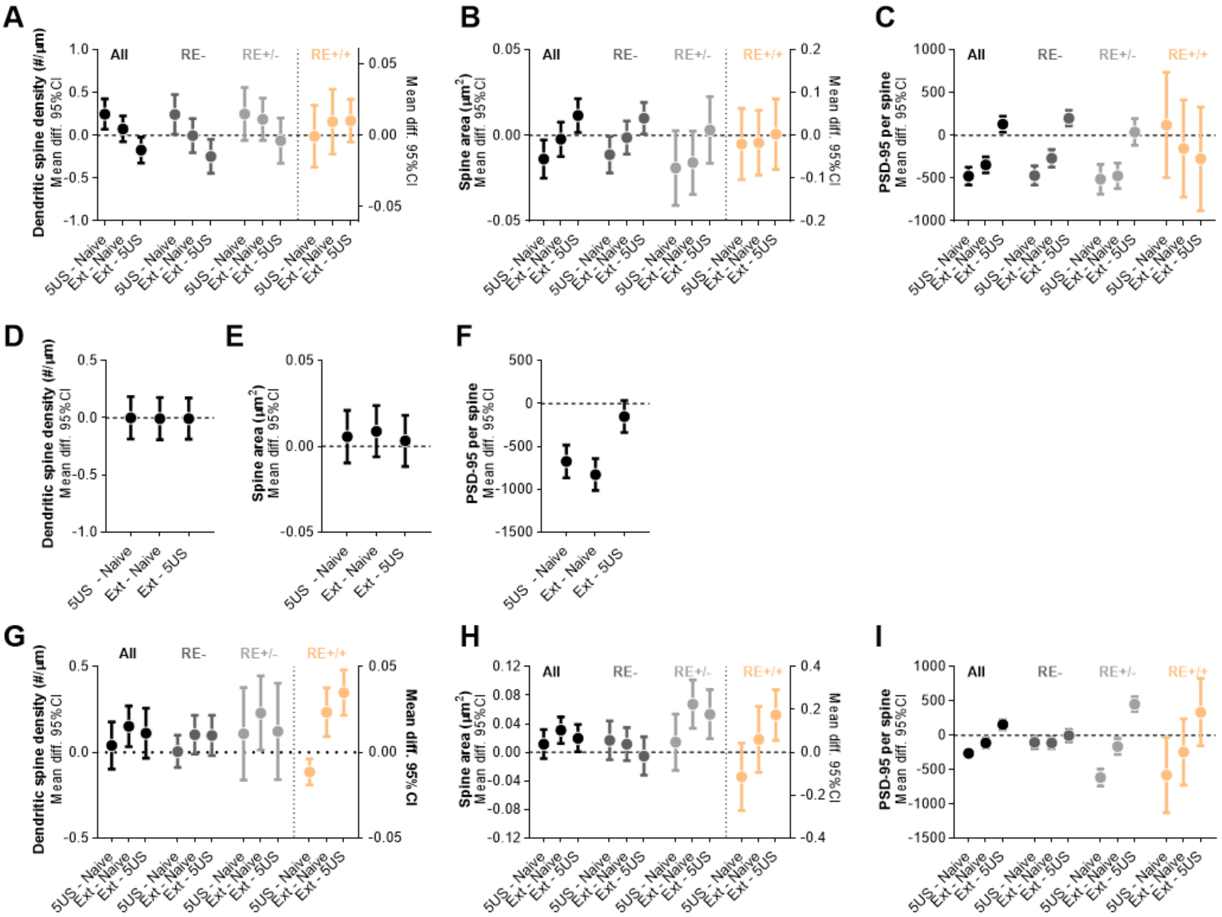


**Supplementary Figure 1. Extinction of contextual fear remodels SLM dendritic spines on RE+ dendrites (estimation statistics).**

**(A-C)** Analysis of dendritic spines in SR. Effect size for training-induced changes of **(A)** dendritic spine density (RE+/+ spines are plotted on the right Y axis), **(B)** dendritic spine area (RE+/+ spines are plotted on the right Y axis) and **(C)** PSD-95 expression per dendritic spine.
**(D-F)** Analysis of dendritic spines in SR. Effect size for training-induced changes of **(D)** dendritic spine density, **(E)** dendritic spine area and **(F)** PSD-95 expression per dendritic spine.
**(G-I)** Analysis of dendritic spines in SR. Effect size for training-induced changes of **(G)** dendritic spine density (RE+/+ spines are plotted on the right Y axis), **(H)** dendritic spine area (RE+/+ spines are plotted on the right Y axis) and **(I)** PSD-95 expression per dendritic spine.

**Supplementary Table 1. 95% CI for training-induced changes in density of dendritic spines in SO.**

| | Total | | | RE- | | | RE+/- | | | RE+/+ | | |
|---|---|---|---|---|---|---|---|---|---|---|---|---|
| | Mean | Upper limit | Lower limit | Mean | Upper limit | Lower limit | Mean | Upper limit | Lower limit | Mean | Uppe r limit | Lower limit |
| 5US - Naive | 0,262 | 0,447 | 0,077 | 0,268 | 0,515 | 0,021 | 0,249 | 0,531 | -0,032 | -0,0006 | 0,281 | -0,282 |
| Ext - Naive | 0,059 | 0,228 | -0,109 | -0,025 | 0,202 | -0,253 | 0,187 | 0,441 | -0,066 | 0,009 | 0,263 | -0,244 |
| Ext - 5US | -0,203 | -0,045 | -0,361 | -0,293 | -0,092 | -0,495 | -0,062 | 0,191 | -0,316 | 0,010 | 0,264 | -0,243 |

**Supplementary Table 2. 95% CI for training-induced changes in dendritic spine area in SO.**

| | Total | | | RE- | | | RE+/- | | | RE+/+ | | |
|---|---|---|---|---|---|---|---|---|---|---|---|---|
| | Mean | Upper limit | Lower limit | Mean | Upper limit | Lower limit | Mean | Upper limit | Lower limit | Mean | Uppe r limit | Lower limit |
| 5US - Naive | -0,0138 | -0,002 | -0,024 | -0,011 | -0,000 6 | -0,022 | -0,019 | 0,0027 | -0,040 | -0,0202 | 0,063 | -0,103 |
| Ext - Naive | -0,0022 | 0,0077 | -0,012 | -0,001 | 0,0082 | -0,011 | -0,015 | 0,0025 | -0,034 | -0,0174 | 0,057 | -0,0928 |
| Ext - 5US | 0,0115 | 0,0213 | 0,0016 | 0,0099 | 0,0191 | 0,0007 | 0,0031 | 0,0225 | -0,016 | 0,0028 | 0,084 | -0,0793 |

**Supplementary Table 3. 95% CI for training-induced changes in PSD-95 protein levels in SO.**

| | Total | | | RE- | | | RE+/- | | | RE+/+ | | |
|---|---|---|---|---|---|---|---|---|---|---|---|---|
| | Mean | Upper limit | Lower limit | Mean | Upper limit | Lower limit | Mean | Upper limit | Lower limit | Mean | Upper limit | Lower limit |
| 5US - Naive | -475 | -370 | -579 | -468 | -357 | -580 | -512 | -338 | -686 | 120 | 733 | -492 |
| Ext - Naive | -344 | -253 | -436 | -268 | -166 | -370 | -472 | -323 | -620 | -154 | 414 | -723 |
| Ext - 5US | 130 | 223 | 36 | 200 | 290 | 109 | 40 | 195 | -115 | -274 | 329 | -878 |

**Supplementary Table 4. 95% CI for training-induced changes in density of dendritic spines in SR.**

| | Total | | |
|---|---|---|---|
| | Mean | Upper limit | Lower limit |
| 5US - Naive | 0,0005 | 0,1854 | -0,184 |
| Ext - Naive | -0,0065 | 0,1782 | -0,191 |
| Ext - 5US | -0,0071 | 0,1730 | -0,187 |

**Supplementary Table 5. 95% CI for training-induced changes in dendritic spine area in SR.**

| | Total | | |
|---|---|---|---|
| | Mean | Upper limit | Lower limit |
| 5US - Naive | 0,0055 | 0,0209 | -0,009 |
| Ext - Naive | 0,0087 | 0,0237 | -0,006 |
| Ext - 5US | 0,0031 | 0,0181 | -0,011 |

**Supplementary Table 6. 95% CI for training-induced changes in PSD-95 protein levels in SR.**

| | Total | | |
|---|---|---|---|
| | Mean | Upper limit | Lower limit |
| 5US - Naive | -676 | -485 | -867 |
| Ext - Naive | -829 | -643 | -1015 |
| Ext - 5US | -152 | 33 | -338 |

**Supplementary Table 7. 95% CI for training-induced changes in density of dendritic spines in SLM.**

| | Total | | | RE- | | | RE+/- | | | RE+/+ | | |
|---|---|---|---|---|---|---|---|---|---|---|---|---|
| | Mean | Upper limit | Lower limit | Mean | Upper limit | Lower limit | Mean | Upper limit | Lower limit | Mean | Upper limit | Lower limit |
| 5US - Naive | 0,039 | 0,177 | -0,097 | 0,0047 | 0,0992 | -0,089 | 0,1077 | 0,3767 | -0,161 | -0,011 4 | -0,003 | -0,0189 |
| Ext - Naive | 0,151 | 0,2697 | 0,0329 | 0,1025 | 0,2160 | -0,010 | 0,2293 | 0,4453 | 0,0133 | 0,0233 | 0,037 | 0,009 |
| Ext - 5US | 0,111 | 0,256 | -0,033 | 0,097 | 0,214 | -0,019 | 0,121 | 0,402 | -0,159 | 0,034 | 0,047 | 0,0215 |

**Supplementary Table 8. 95% CI for training-induced changes in dendritic spine area in SLM.**

| | Total | | | RE- | | | RE+/- | | | RE+/+ | | |
|---|---|---|---|---|---|---|---|---|---|---|---|---|
| | Mean | Upper limit | Lower limit | Mean | Upper limit | Lower limit | Mean | Upper limit | Lower limit | Mean | Uppe r limit | Lower limit |
| 5US - Naive | 0,011 | 0,0316 | -0,008 | 0,016 | 0,043 | -0,010 | 0,014 | 0,053 | -0,025 | -0,114 | 0,042 | -0,270 |
| Ext - Naive | 0,031 | 0,049 | 0,012 | 0,011 | 0,034 | -0,011 | 0,0673 | 0,101 | 0,033 | 0,060 | 0,213 | -0,093 |
| Ext - 5US | 0,019 | 0,038 | 0,0005 | -0,005 | 0,0217 | -0,032 | 0,0531 | 0,0875 | 0,0187 | 0,174 | 0,292 | 0,055 |

**Supplementary Table 9. 95% CI for training-induced changes in PSD-95 protein levels in SLM.**

| | Total | | | RE- | | | RE+/- | | | RE+/+ | | |
|---|---|---|---|---|---|---|---|---|---|---|---|---|
| | Mean | Upper limit | Lower limit | Mean | Upper limit | Lower limit | Mean | Upper limit | Lower limit | Mean | Upper limit | Lower limit |
| 5US - Naive | -265 | -192 | -337 | -105 | -17 | -192 | -617 | -490 | -744 | -579 | -30 | -1129 |
| Ext - Naive | -111 | -44 | -179 | -111 | -29 | -193 | -165 | -49 | -281 | -245 | 236 | -727 |
| Ext - 5US | 153 | 222 | 84 | -5 | 80 | -92 | 452 | 561 | 344 | 334 | 826 | -158 |
